# FeLIX is a restriction factor for mammalian retrovirus infection

**DOI:** 10.1101/2023.11.14.567074

**Authors:** Didik Pramono, Dai Takeuchi, Masato Katsuki, Loai AbuEed, Dimas Abdillah, Toru Kimura, Junna Kawasaki, Ariko Miyake, Kazuo Nishigaki

## Abstract

Endogenous retroviruses (ERVs) are remnants of ancestral viral infections. Feline leukemia virus (FeLV) is an exogenous and endogenous retrovirus in domestic cats. It is classified into several subgroups (A, B, C, D, E, and T) based on viral receptor interference properties or receptor usage. ERV-derived molecules benefit animals, conferring resistance to infectious diseases. However, the soluble protein encoded by the defective envelope (*env*) gene of endogenous FeLV (enFeLV) functions as a co-factor in FeLV subgroup T infections. Thus, whether the gene emerged to facilitate viral infection is unclear. Based on the properties of ERV-derived molecules, we hypothesized that the defective *env* genes possess antiviral activity that would be advantageous to the host because FeLV subgroup B (FeLV-B), a recombinant virus derived from enFeLV *env*, is restricted to viral transmission among domestic cats. When soluble truncated Env proteins from enFeLV were tested for their inhibitory effects against enFeLV and FeLV-B, they inhibited viral infection. Notably, this antiviral machinery was extended to infection with the Gibbon ape leukemia virus, Koala retrovirus-A, and Hervey pteropid gammaretrovirus. Although these viruses used feline phosphate transporter1 (fePit1) or fePit1 and phosphate transporter2 (fePit2) as receptors, the inhibitory mechanism involved competitive receptor binding in a fePit1-dependent manner. The shift of receptor usage may have occurred to avoid the inhibitory effect. Overall, these findings highlight the possible emergence of soluble truncated Env proteins from enFeLV as a restriction factor against retroviral infection, and might help in the control of retroviral spread for host immunity and antiviral defense.

**Importance:** Retroviruses are unique in using reverse transcriptase to convert RNA genomes into DNA, infecting germ cells, and transmitting to offspring. A large amount of ancient retroviral sequences are known as endogenous retroviruses (ERVs). Soluble Env protein derived from ERVs have been identified to function as a co-factor that assists in FeLV-T infection. However, herein, we show that the soluble Env protein exhibits antiviral activity and provides resistance to mammalian retrovirus infection through competitive receptor binding. In particular, this finding may explain why FeLV-B transmission is not observed among domestic cats. ERV-derived molecules can benefit animals in an evolutionary arm race, highlighting the double-edged sword nature of ERVs.

## INTRODUCTION

Retroviruses are RNA viruses that infect a wide range of vertebrate hosts, including mammals (1). Retroviruses are unique because they use reverse transcriptase to convert their RNA genome into DNA, which can then be integrated into the host genome. Enormous amounts of ancient retroviral sequences known as endogenous retroviruses (ERVs) infect germ cells, integrate into the host genome, and are vertically transmitted to offspring (2-4). Some ERV Envelope (Env) proteins may have a potent evolutionary significance in the immunological response by acting as restriction factors against exogenous retroviruses (5, 6). Existing research recognized the critical role played by the *env* gene derived from ERVs (env-ERV), which reportedly exhibits restrictive activity against retroviruses in mice (Fv4, Rmcf, and Rmcf2), sheep (enjS56A1), cats (Refrex-1), and humans (Suppressyn) (5-9). The antiviral activity of env-ERV genes is likely owing to their ability to interfere with the entry of exogenous retroviruses into host cells (5-7, 10, 11). The retroviral envelope binds to receptors on the host cell surface and mediates cell entry. Env-ERV proteins compete with exogenous retroviral Env proteins for binding to these receptors, thereby blocking viral entry and replication (5, 7).

Feline leukemia virus (FeLV), a gammaretrovirus endemic to domestic cats worldwide, was identified as the infectious agent responsible for feline leukemia and lymphoma in the early 1960s (12). FeLV induces malignant hematopoietic disorders, including lymphoma, myelodysplastic syndrome (MDS), acute myeloid leukemia (AML), aplastic anemia, and immunodeficiency in cats (13, 14). FeLVs can be categorized into several subgroups based on viral receptor interference and host range. FeLV subgroups and their FeLV-A, -B, -C, -D, -E, and -T receptors have been identified (5, 15-24). FeLV-B arises through recombination in the *env* region between FeLV-A and the endogenous FeLV (enFeLV) present in the feline genome (25-27). FeLV-A is the primary virus transmitted horizontally among domestic cats. Although horizontal transmission of FeLV-B mostly does not occur, it may rarely be transmitted by coinfection with FeLV-A (27). Given that *in vivo* experimental infections have shown resistance to FeLV-B (28), domestic cats are resistant to FeLV-B infection; however, the underlying mechanism is unknown. FeLV-B uses phosphate transporter 1 (Pit1) and phosphate transporter 2 (Pit2) as entry receptors (17, 29). FeLV-B occurs in 33–68% of cats infected with FeLV-A, presumably by independent recombination events that occur *de novo* after infection of domestic cats with FeLV-A (12, 17, 27). In addition to FeLV-B, various other viruses such as KoRV, Gibbon ape leukemia virus (GaLV), Hervey pteropid gammaretrovirus (HPG), 4070A MuLV, 10A1 MuLV, and woolly monkey virus use phosphate transporters as viral receptors. They can spread to hosts, and interspecies transmission can occur (30).

FeLV-T is a variant of FeLV with a mutation in the PHQ motif of Env (16, 31). FeLV-T utilizes Pit-1 as a receptor but lacks inherent cell fusion ability (32) and requires an assisting molecule, namely FeLIX, for viral infection (16). FeLIX is a soluble, truncated Env protein derived from enFeLV, which is released from cells and is present in the culture supernatant of feline cells (16) and serum from domestic cats (33). FeLV-T is not known to exist as a replication-competent virus, but its replication is supported by helper viruses such as FeLV-A. FeLV-T is rarely found in natural cases (25, 26, 34). Of note, experimental inoculation of a recombinant replication-competent FeLV-T showed severe immunodeficiency in domestic cats (35).

FeLV-B transmission does not occur among domestic cats, supporting that *in vivo* experimental infection shows resistance to FeLV-B infection (28). The presence of enFeLV has been suggested to be associated with this phenomenon, and FeLV-B infection in Florida panthers lacking enFeLV highlights the association between enFeLV and FeLV-B (36). The presence of ERV is known to confer resistance to exogenous retroviral infection (5-9, 37, 38). In particular, the recently identified soluble truncated Env proteins confer resistance to viral infection of FeLV-D or RD-114 (5, 6); however, the molecular mechanism by which FeLV-B infection is restricted in domestic cats has long remained unexplained. FeLIX has been identified as a co-factor for FeLV-T infection (16) before the possibility of antiviral activity of FeLIX was investigated. The idea that FeLIX has evolutionarily emerged to facilitate viral infection seems unlikely because it has a detrimental function in domestic cats. Therefore, this study aimed to explore the antiviral activities of FeLIX and enFeLV-derived truncated Env.

## RESULTS

### Genetic diversity of enFeLV in domestic cats

Much remains unknown regarding the interaction between FeLV-B and enFeLV, and complete analyses of enFeLV in domestic cats have not been conducted. Therefore, in this study, we used two strategies to identify enFeLV sequences in domestic cats. The first approach involved searching for publicly available whole-genome sequence data from domestic cats. We downloaded all available domestic cat sequence data using the *env* gene of enFeLV-AGTT as a reference because our objective was to determine the Env function of enFeLV. Our second approach involved genomic library screening and PCR using locus-specific primers. Sequence analysis revealed six proviruses carrying the *env* gene, namely enFeLV-clone1, -clone2, -clone3, -clone4, -clone5, and -clone6; three of these proviruses (enFeLV-clone1, enFeLV-clone2, and enFeLV-clone3) had intact open reading frames (ORFs) for *env* (Figure S1), whereas the other three (enFeLV- clone4, enFeLV-clone5, and enFeLV-clone6) had defective *env* genes. Sequence analysis of the database identified that enFeLV-clone5, referred to as CFE16 (39) and FeLIX (16), carried a truncated *env* gene. The truncated enFeLV-clone4 Env was termed Trunc-C4. EnFeLV-clone6 had an ORF as the defective *env* gene. The obtained proviruses data (Table S1) were comprehensively analyzed via the phylogenetic tree using 5ʹ and 3ʹ long terminal repeat (LTR) sequences, where enFeLV grouped differently from exogenous FeLV (exFeLV) (Figure 1A). Examination of the chromosomal location of enFeLV in the cat genome revealed that it was widely present on almost all chromosomes, with chromosome B4 having the highest copy number; however, no enFeLV was found on chromosomes C1, C2, D2, E1, E2, E3, and F2 (Figure 1B). The 5ʹ and 3ʹ LTR sequences of each enFeLV were found to have high sequence identity or were identical to each other. The integration time of each enFeLV locus was estimated by comparing the nucleotide mismatch between the 5ʹ and 3ʹ LTRs (Table S2). EnFeLVs carrying *env* were estimated to have been endogenized less than 1 million years ago (Figure 1C). Notably, enFeLV-clone5 (CFE-16 and FeLIX) and enFeLV-clone4 were endogenized less than 200 thousand years ago (Table S2).

**Figure 1.**
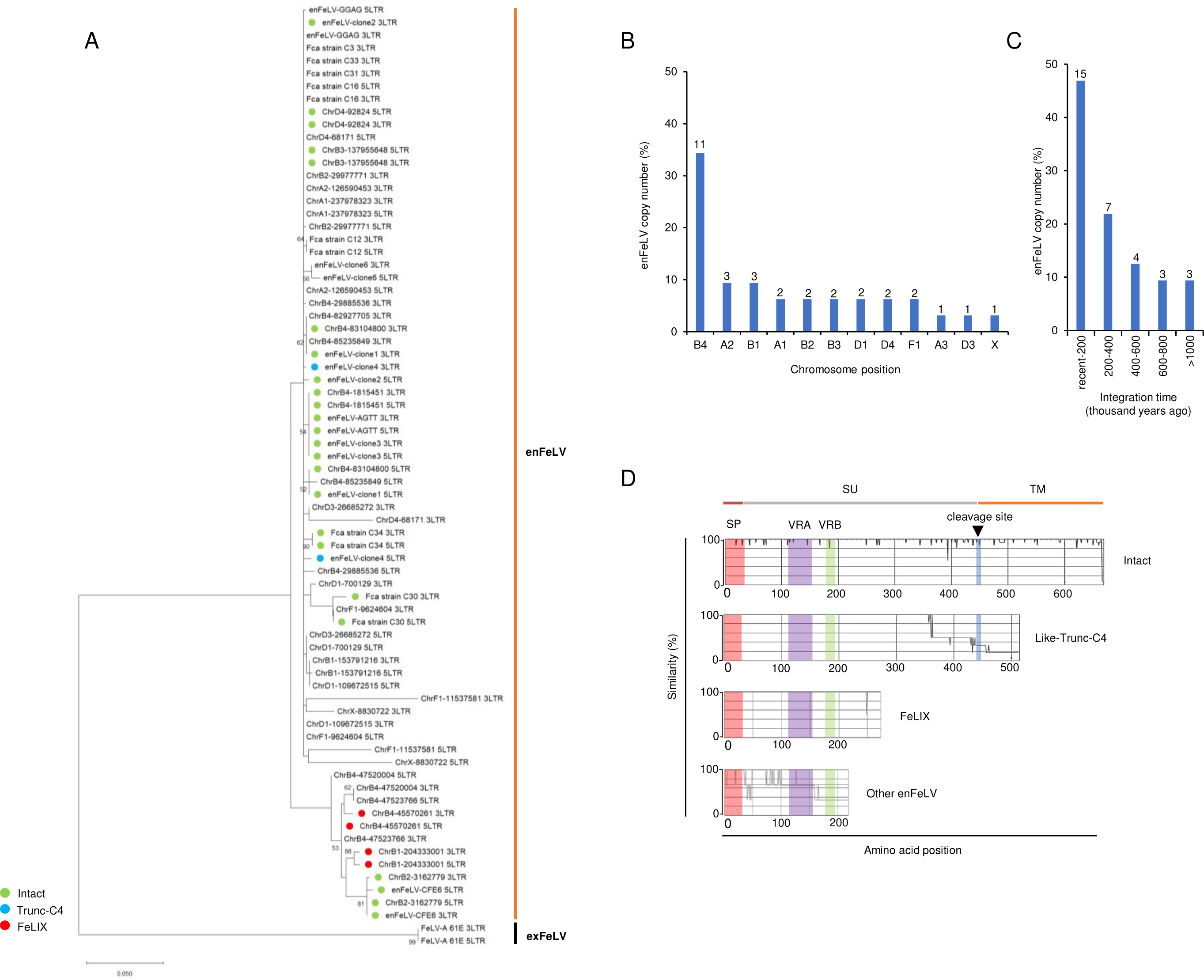
Endogenization of enFeLV in domestic cats. (A) Phylogenetic tree of enFeLV carrying the *env* gene based on LTRs. FeLIX and Trunc-C4 are marked by red and blue circles, respectively. (B) Chromosome position of enFeLV proviral integration in domestic cat genome. The number of viral integrations is indicated in the top bar. (C) Integration time of enFeLV proviruses in domestic cat genome. (D) Structure of enFeLV Env classification (4 groups); Group 1: Intact, Intact full-length Env has approximately 666 amino acids (aa); Group 2: Like-Trunc-C4, defective Env like-Trunc-C4 (length > 300 aa), which has SP (Signal peptide), VRA (Variable region A), and VRB (Variable region B) in Env SU (Surface Unit) but lacks TM (Transmembrane domain) region; Group 3: like-FeLIX (length < 300 aa), which has only SP, VRA, and VRB in Env SU; Group 4; Env has a length between 140–273 aa and lacks VRA, VRB, or the TM region.

Next, we divided the enFeLV Env structures into four groups (Figures 1D and S1). The first group, intact full-length Env, had approximately 666 amino acids with a signal peptide (SP), surface unit (SU) containing variable region A (VRA) and variable region B (VRB), receptor-binding domain (RBD), and transmembrane (TM) region. The second group, defective Env like-Trunc-C4 (length > 300 aa), had SP, VRA, and VRB in Env SU but lacked the TM region. The third group, FeLIX (CFE16, enFeLV-clone 4; length = 273 aa), had only SP, VRA, and VRB in Env SU, and the fourth group, Env, had a length of 140–273 aa and lacked the VRA, VRB, or TM region. Taken together, the structural diversity of Env was generated less than 1 million years ago (Mya) (Figure S1).

### Infectivity of enFeLV Env-pseudotyped virus

To determine whether the intact *env* genes of enFeLVs form infectious viral particles using MLV Gag-Pol and MuLV retroviral vectors carrying LacZ as a marker, GPLac cells were transfected with the *env* gene to produce the enFeLV Env-pseudotyped virus. The results showed that enFeLV-clone2, enFeLV-clone3, and enFeLV-AGTT infected AH927 and CRFK cells as the target cells (Figure 2A). They showed high viral titers comparable to those observed in FeLV-B infection (Figure 2A). In contrast, enFeLV- clone1 did not cause infection. The expression of enFeLV-clone1 Env, as demonstrated via western blot analysis, showed incomplete cleavage between SU (gp70) and TM (p15e; Figure 2C). Amino acids involved in infectivity were determined by comparing the amino acid sequences of enFeLV-clone1 and enFeLV Env (clone2 and clone3). The enFeLV-clone1 mutants E345G (substitution of glutamic acid with glycine at position 345) and N394K (substitution of asparagine with lysine at position 394) were tested for infection with their Env-pseudotyped viruses. The enFeLV-clone1 E345G showed infectivity (Figure 2B), and the SU-TM region was cleaved into Env (Figure 2C). In contrast, the other mutation (N394K) did not infect target cells (Figure 2B) and was not cleaved by Env (Figure 2C). This confirmed that the amino acid change from G (glycine) to E (glutamic acid) at position 345 caused cleavage failure of the Env protein, resulting in the loss of viral infectivity. These results suggest that enFeLV Env can generate infectious particles that may pose a threat to domestic cats.

**Figure 2.**
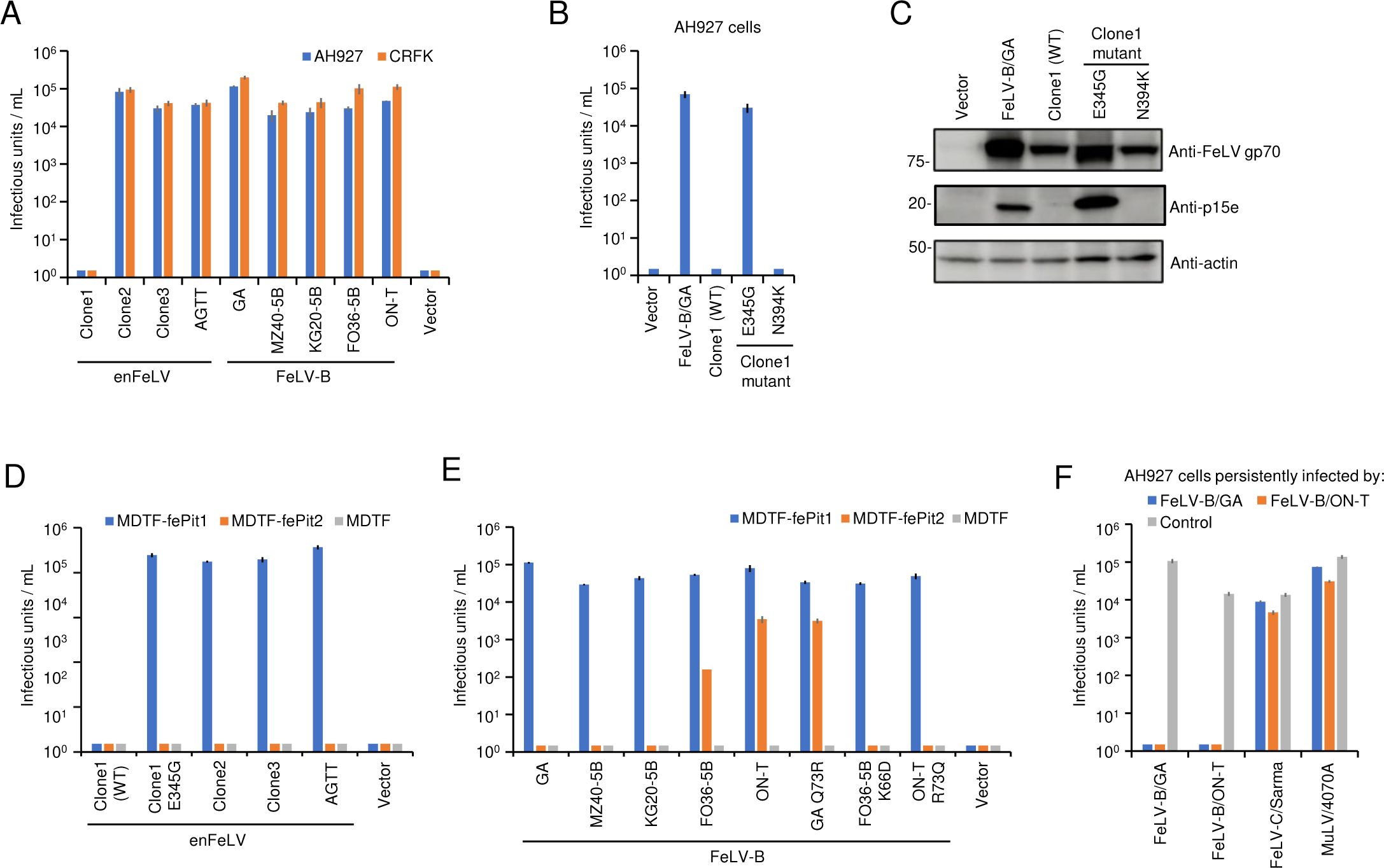
Infectivity and receptor usage of enFeLV Env-pseudotyped viruses. (A) Infectivity of enFeLV (clone1, clone2, and clone3) and FeLV-B (GA, MZ40-5B, KG20- 5B, FO36-5B, and ON-T) Env-pseudotyped viruses. (B) Infectivity of Env mutants (E345G and N394K) from enFeLV clone1. (C) Western blot analysis of enFeLV Envs proteins in HEK293T cells using anti-FeLV gp70 and anti-FeLV p15e antibodies. Actin was used as a control. (D) Infection assay of enFeLV (clone1/WT, clone1 E345G, clone2, clone3, and AGTT) in MDTF-fePit1, MDTF-fePit2, and MDTF (empty vector) as target cells for receptor usage. (E) Infection assay of FeLV-B (GA, MZ40-5B, KG20- 5B, FO36-5B, and ON-T) and FeLV-B mutants (GA Q73R, FO36-5B K66D, and ON-T R73Q) in MDTF-fePit1, MDTF-fePit2, and MDTF (empty vector) as target cells for receptor usage. (F) Interference assay of FeLV-B/GA and FeLV-B/ON-T. AH927 cells pre-infected with either FeLV-B/GA or FeLV-B/ON-T were infected by the Env-pseudotyped viruses. FeLV-C/Sarma and MuLV 4070A were used as controls. Viral titers are indicated on the x-axis. The infectious units (IU) were determined by counting the number of log_10_-galactosidase (LacZ)-positive cells per milliliter (mL) of virus indicated on the y-axis. Virus infection titers with standard deviations were averaged from three independent experiments. Mock represents the negative control.

### Identification of an entry receptor for enFeLV

FeLV-B recombines in the *env* region between FeLV-A and enFeLV (25-27). We hypothesized that enFeLV used the same receptor as FeLV-B, termed feline Pit1 (fePit1) and feline Pit2 (fePit2). fePit1 and fePit2 are members of the family of solute carrier proteins SLC20A1 and SLCA20A2, structurally identical sodium-dependent phosphate transporters. *Mus dunni* tail fibroblast (MDTF) cells that were resistant to enFeLV infection (Figure 2D), MDTF cells expressing fePit1 (MDTF-fePit1), and MDTF cells expressing fePit2 (MDTF-fePit2) were tested for infection with enFeLV Env-pseudotyped viruses prior to analysis 2 days later. The results showed that MDTF-fePit1 cells were permissive to enFeLV Env-pseudotyped virus infection of clone1 E345G, clone2, clone3, and AGTT, whereas MDTF-fePit2 cells were not infected with any of the enFeLV Env-pseudotyped viruses (Figure 2D). Interestingly, both enFeLV-clone1, estimated to have integrated approximately 220 thousand years ago, and clone2, a recent addition, utilize the same receptor (Table S3).

Next, we evaluated the receptor usage of the five FeLV-B strains, showing the structural diversity of *env* genes by recombination analysis (26) (Figure S2). MDTF-fePit1 cells were permissible for infection with all FeLV-B Env-pseudotyped viruses (GA, MZ40-5B, KG20-5B, FO36-5B, and ON-T), whereas only two FeLV-B Env-pseudotyped viruses (FO36-5B and ON-T) could infect MDTF-fePit2 cells (Figure 2E). Based on the amino acid sequence of FeLV-B *env*, lysine at amino acid position 66 and arginine at position 73 in *env* variable region A (VRA) may be associated with a shift in receptor use. Therefore, FeLV-B/FO36-5B K66D (lysine to aspartic acid) and FeLV-B/ON-T R73Q (arginine to glutamine) mutants were tested for receptor usage. FeLV-B/FO36-5B K66D and FeLV-B/ON-T R73Q infected MDTF-fePit1 cells but not MDTF-fePit2 cells. Furthermore, FeLV-B/GA Q73R (glutamine to arginine) infected both MDTF-fePit1 and MDTF-fePit2 cells, whereas FeLV-B/GA only infected MDTF-fePit1 cells (Figure 2E). These results indicate that lysine at position 66 and arginine and glutamine at position 73 affect the receptor usage of FeLV-B infection. We conducted an interference assay with FeLV-B/GA (fePit1) and FeLV-B/ON-T (fePit1 and fePit2) in AH927 cells. The pseudotyped viruses of FeLV-B/GA did not infect the AH927 cells infected with FeLV-B/ON-T. Conversely, FeLV-B/ON-T did not infect the AH927 cells infected with FeLV-B/GA (Figure 2F). These results suggested that blocking the fePit1-mediated infection pathway was sufficient to prevent FeLV-B infection. In other words, the major pathway for FeLV-B infection may involve fePit1.

### Expression of fePit1 and fePit2 in feline tissues and cell lines

Proteins of 681 and 653 amino acids were predicted to be encoded by fePit1 and fePit2, respectively (17, 40). Approximately 88% of the amino acids in fePit1 and fePit2 are identical. qRT-PCR showed the expression of fePit1 and fePit2 mRNA; the cerebellum, large intestine, ovary, and spleen exhibited the highest levels of fePit1 expression. The cerebellum, kidney, uterus, and liver exhibited the highest fePit2 expression (Figure S3A). The fePit1 and fePit2 genes were detected in all feline cell lines, including AH927, CRFK, Fet-J, MCC, MS4, and 3201 (Figure S3B). Hematopoietic cell lines tended to express pit1 and pit2. These findings indicate that fePit1 and fePit2 are broadly expressed in feline tissue and cell lines.

### Truncated Envs from enFeLV

FeLIX (CFE16, enFeLV-clone5) and Trunc-C4 had similar structures, with lengths of 273 aa and 369 aa, respectively (Figure 3A). We examined whether truncated *env* genes expanded across domestic cats in Japan. Proviruses encoding FeLIX on chromosomes B1 (Chr B1) and B4 were detected using PCR in all domestic cats (Figure 3B). In addition to examining the loci, a single-nucleotide polymorphism (SNP) in the genes encoding FeLIX was identified as either aspartic acid (D) or asparagine (N) at amino acid position 249 on chromosome B1. Asparagine (N) was found at amino acid position 249 in B4. These cells were termed FeLIX-N249 and FeLIX-D249, respectively. Notably, an SNP of FeLIX was located within the FeLV-A-neutralizing epitope “MGPNL” at amino acid positions 246–250 in FeLV-A/61E Env. Furthermore, the neutralizing epitope was not conserved in FeLIX-N249 (MGPNP) and FeLIX-D249 (MGPDP) and contained a substitution of proline (Figure S4). Trunc-C4 was detected in 12 of 22 cats, suggesting that Trunc-C4 is not fixed in domestic cats (Figure S5). We determined whether FeLIX and Trunc-C4 are evolutionarily conserved in felids. We detected FeLIX-D249 on chromosome B1 and FeLIX-N249 on chromosome B4 in all European wild cats (nine cats); however, Trunc-C4 was not detected. Since the 5ʹ LTR and 3ʹ LTR of Trunc-C4 showed a high genetic distance between them, this provirus could be generated by recombination; therefore, the integration time calculated by LTRs may not support that estimated based on animal segregation (Figure 1A). These results suggest that FeLIX emerged as a common ancestor of domestic cats and European wild cats.

**Figure 3.**
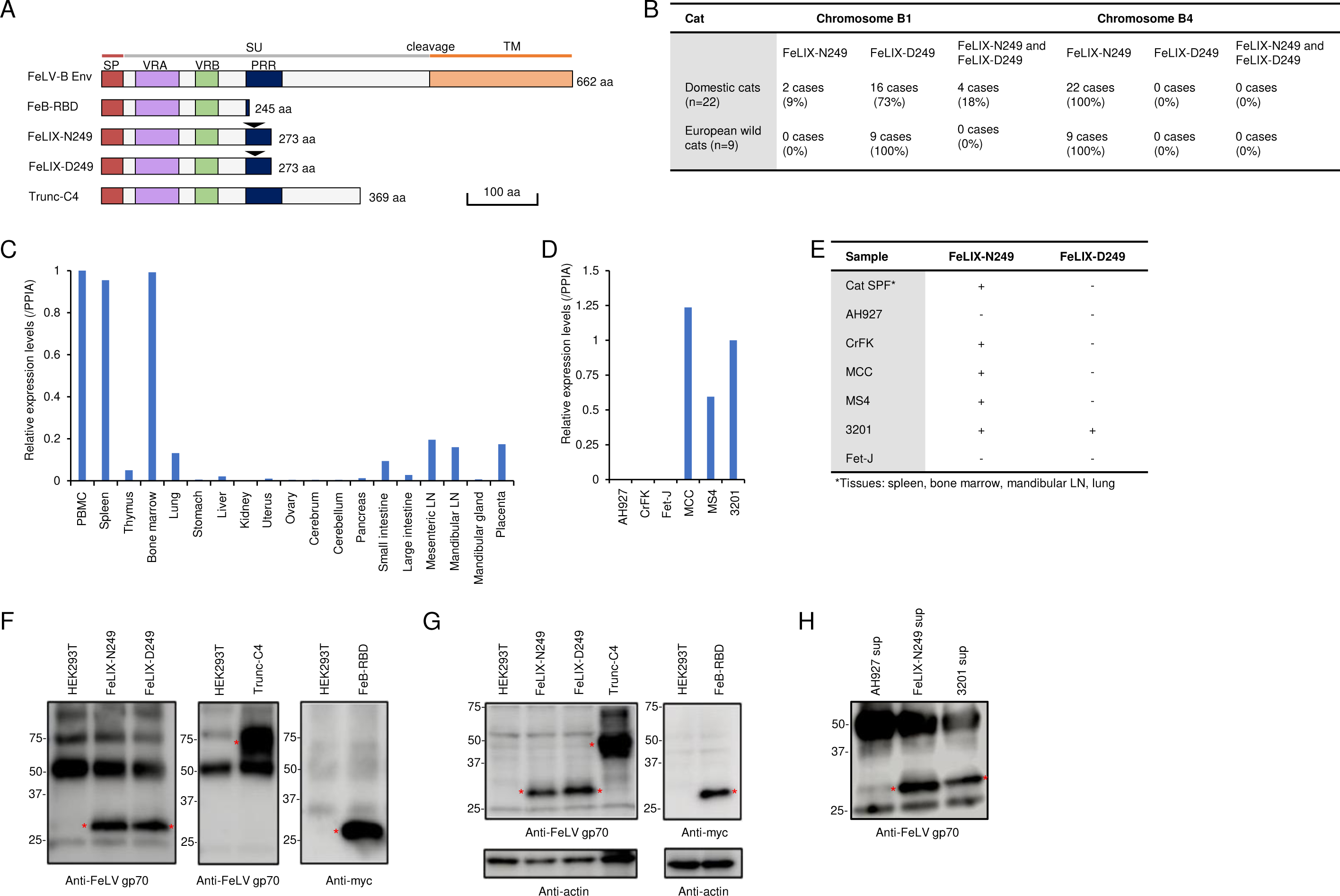
Structural schematic and expression of truncated Envs. (A) Schematic representation of the structure of FeLIX-N249 (asparagine at position 249), FeLIX-D249 (aspartic acid at position 249), and Trunc-C4. The receptor-binding domain (RBD) of FeLV-B/GA with myc-tag is also shown. SP, signal peptide; SU, surface unit; TM, transmembrane domain; VRA, variable region A; VRB, variable region B; PRR, proline-rich region. The number of amino acids is indicated on the right side. (B) Detection for proviruses encoding FeLIX in the domestic cat (N-22) and European wild cat (n = 9) genome. (C, D) FeLIX expression in feline tissues and cell lines. Quantification of feline FeLIX transcripts by quantitative RT-PCR in feline tissues and cell lines. The x-axis indicates the analyzed samples. The y-axis indicates the expression level normalized to the expression of peptidylprolyl isomerase A (PPIA). Normalized expression in PBMC and 3201 cells is shown as 1 in feline tissues and cell lines, respectively. LN, Lymph node. (E) Expression of FeLIX-N249 and D-249 was determined via RT-PCR and sequencing in feline tissues (spleen, bone marrow, mandibular LN, and lung) and indicated feline cell lines. (F) Detection of FeLIX-N249 and D-249, Trunc-C4, and Fe-B-RBD in cell culture supernatants and (G) in cell lysates from HEK293T transfected with indicated plasmids. (H) Detection of FeLIX in supernatant of indicated cells. Red asterisks indicate truncated Env proteins. FeLIX-N249 and D-249, as well as Trunc-C4, were analyzed via immunoprecipitation (IP) and western blotting (WB) using a goat anti-FeLV gp70 antibody (81S-210-2, NCI), while Fe-B-RBD was assessed with an anti-Myc antibody.

Next, we quantified FeLIX expression in normal feline tissues (SPF cat) and cell lines using qRT-PCR. FeLIX is broadly expressed in tissues, with the highest levels identified in immune system tissues (spleen, bone marrow, and lymph nodes) and PBMCs. FeLIX was detected in 3201 (T-cell lymphoma), MS4 (B-cell lymphoma), and MCC (large granular lymphoma) cells (Figure 3C and 3D). AH927, CRFK (kidney cells), and Fet-J cells were barely detectable. Sequence analysis showed that both FeLIX-N249 and FeLIX-D249 were expressed in 3201 cells, whereas only FeLIX-N249 was expressed in all other cells as well as in tissues from the spleen, bone marrow, mandibular lymph node, and lungs from one cat (Figure 3E). We could not determine the expression of Trunc-C4 in domestic cats because of the high sequence similarity with the enFeLV sequences. Next, we determined the expression of FeLIX and Trunc-C4 as soluble proteins. The supernatants of HEK293T cells transfected with FeLIX-N249, FeLIX- D249, and Trunc-C4 expression plasmids were immunoprecipitated using an anti-FeLV gp70 antibody, and western blot analysis was conducted using the same antibody (Figure 3F and 3G). The results showed that these proteins were present in the cell supernatants, suggesting that FeLIX-N249, FeLIX-D249, or Trunc-C4 was secreted from cells as soluble proteins.

### Truncated Env proteins derived from enFeLV confer resistance to enFeLV and FeLV-B infection

FeLIX acts as a co-factor for FeLV-T infection (16). However, truncated ERV Env proteins exhibit antiviral functions (6, 7). Therefore, we investigated whether truncated Env proteins from enFeLVs showed antiviral activity, which benefits the host. HEK293T cells were transfected with each expression plasmid, and the supernatants were collected as sources of FeLIX-N249, FeLIX-D249, and Trunc-C4 proteins. The RBD from FeLV-B/GA, termed FeB-RBD, was used in this study because it showed an inhibitory effect against the FeLV-B/GA strain (41). Next, FeLIX-N249, FeLIX-D249, Trunc-C4, and FeB-RBD were assessed for their inhibitory effects on enFeLV Env-pseudotyped viruses (clone1 E345G, clone2, clone3, and AGTT) and FeLV-B (GA, MZ40-5B, KG20-5B, FO36-5B, and ON-T) in AH927 and CRFK cells. The target cells were treated with the truncated Env proteins for 2 h. Subsequently, pseudotyped viruses were infected into target cells for infection analysis 2 days later. FeLIX, Trunc-C4, and FeB-RBD significantly (*p <* 0.01) inhibited infection with all four enFeLV Env-pseudotyped viruses and all five FeLV-B Env-pseudotyped viruses (Figures 4A, 4B, S6A, S6B, and S7). Notably, different inhibitory effects on enFeLV-clone2 and clone3 were observed between FeLIX-N249 and FeLIX-D249 when HEK293T cells were used as the target cells for the assay (Figure S8). To determine the specificity of the inhibitory effects of FeLIX, Trunc-C4, and FeB-RBD, the inhibition assay was extended to FeLV-A, FeLV-C, and FeLV-E Env-pseudotyped viruses. The results showed that they had no inhibitory effects on FeLV-A, FeLV-C, or FeLV-E infection (Figure 4B). Furthermore, infection of AH927 and CRFK cells by enFeLV-AGTT, FeLV-B/ON-T, and FeLV-B/GA Env-pseudotyped viruses was inhibited in a dose-dependent manner (Figure 4C and S6C). The inhibitory effect of FeLIX on replicative viral infections was tested. A chimeric replication-competent virus carrying enFeLV-AGTT *env* (61E/AGTT) was constructed from FeLV-A/61E. We evaluated FeLIX against the replication-competent infectious viruses FeLV-B/ON-T, FeLV-B/GA, 61E/AGTT, and FeLV-A/61E. Interestingly, FeLIX showed a complete inhibitory effect on replication-competent infectious viruses (Figure 4D and S6D). However, this effect was not observed in FeLV-A 61E infections. Collectively, FeLIX blocked the infection and replication of FeLV-B and enFeLV.

**Figure 4.**
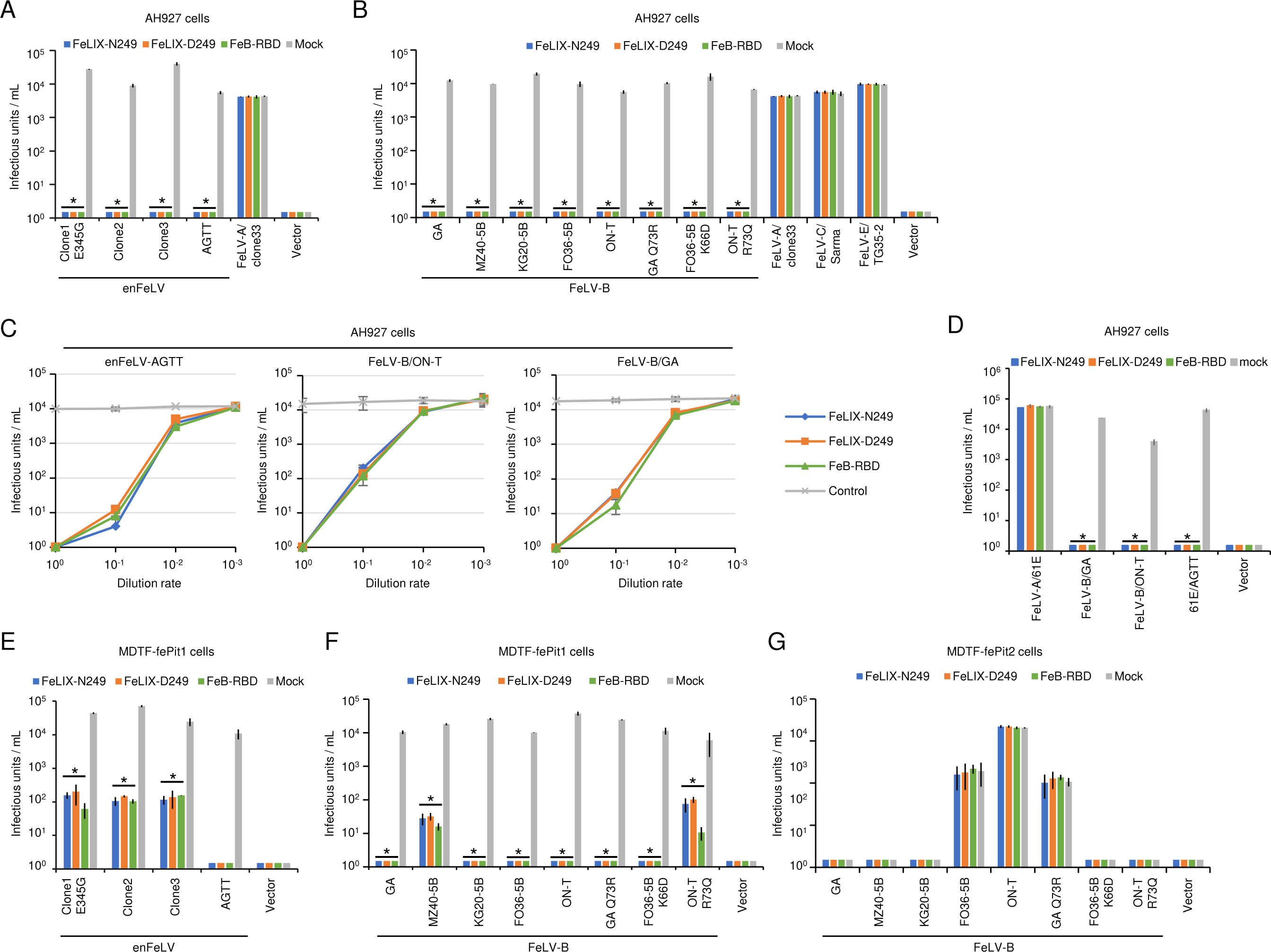
Inhibitory effect of the truncated Env proteins derived from enFeLV for enFeLV and FeLV-B infection and restriction mechanism. (A) Inhibition assay using FeLIX-N249 and FeLIX-D249 for evaluating the infection of Env-pseudotyped viruses, enFeLV (Clone 1 E345G, Clone 2, Clone3, and AGTT), (B) FeLV-B (GA, MZ40-5B, KG20-5B, FO36-5B, and ON-T), and FeLV-B mutants (GA Q73R, FO36-5B K66D, and ON-T R73Q) in AH927 cells. (C) Dose-dependent inhibition of FeLIX for Env-pseudotyped viral infection (FeLV-B/ON-T, FeLV-B/GA, and enFeLV-AGTT) in AH927 cells. (D) against replication-competent viruses assessed included FeLV-B/GA, FeLV-B/ON-T, FeLV-A carrying the enFeLV-AGTT *env* gene, and FeLV-A/61E in AH927 cells. Restriction mechanism in (E, F) MDTF-fePit1 and (G) MDTF-fePit2 cells. FeLV-A/clone33, FeLV-C/Sarma, and FeLV-E/TG35-2 were also used in this assay. FeLIX-N249, FeLIX-D249, FeB-RBD, and the empty vector/mock were sourced from supernatants of HEK293T cells transfected with their respective expression vectors. Each supernatant was added to the culture for 2 h. Subsequently, cells were infected with the Env-pseudotyped virus. The infectious units (IU) shown on the x-axis were determined by counting the number of log_10_-galactosidase (LacZ)-positive cells per milliliter (mL) of virus indicated on the y-axis. Virus infection titers with standard deviations represent the means of three independent infection experiments. Comparisons were performed using Student’s t-test (\**p* < 0.01).

### Feline Pit1-dependent inhibitory effect of FeLIX

We attempted to elucidate the mechanism underlying the inhibitory effect of FeLIX on FeLV-B and enFeLV infection in MDTF-fePit1 and MDTF-fePit2 cells. FeLIX inhibited both retroviral infections in MDTF-fePit1 cells (Figure 4E and 4F). However, FeLV-B/FO36-5B, FeLV-B/ON-T, and FeLV-B/GA Q73R mutants, which showed shifted receptor usage, were not inhibited by FeLIX in MDTF-fePit2 cells (Figure 4G). FeLV-B/ON-T R73Q, FeLV-B/GA Q73R, and FO36-5B K66D, which shift receptor usage, were inhibited by FeLIX in AH927, CRFK, and MDTF-fePit1 cells (Figures 4B, S6B, and 4F). Inhibition by FeB-RBD was fePit1-dependent but not fePit2-dependent, resulting in an effect similar to that of FeLIX. These results suggest that FeLIX inhibits viral infection via fePit1, but not fePit2.

### Inhibitory effects of FeLIX from the supernatant of 3201 cells against FeLV-B and enFeLV infection

FeLIX was present in the supernatants of 3201 cells, as previously reported (16), and qRT-PCR and western blot analyses confirmed the expression of FeLIX in 3201 cells (Figure 3D and 3H). We tested the supernatant of 3201 cells for pseudotyped FeLV-B and enFeLV in AH927 cells as target cells. The supernatant of 3201 cells significantly inhibited infection with FeLV-B (GA, MZ40-5B, KG20-5B, and ON-T) and enFeLV (AGTT, clone 1 E345G, and clone2; *p <* 0.01; Figure 5A). The level of inhibition of FeLV-B GA from the supernatant of 3201 cells tended to be lower than that of other viruses. Infection with FeLV-A, FeLV-C, or FeLV-E was not inhibited in the supernatant of the 3201 cells (Figure 5A). Notably, the antiviral effect of the culture supernatant derived from 3201 cells was not significant when a high viral titer (10^5^ units/mL of FeLV-B/GA) was used to examine the antiviral effect (Figure S9).

**Figure 5.**
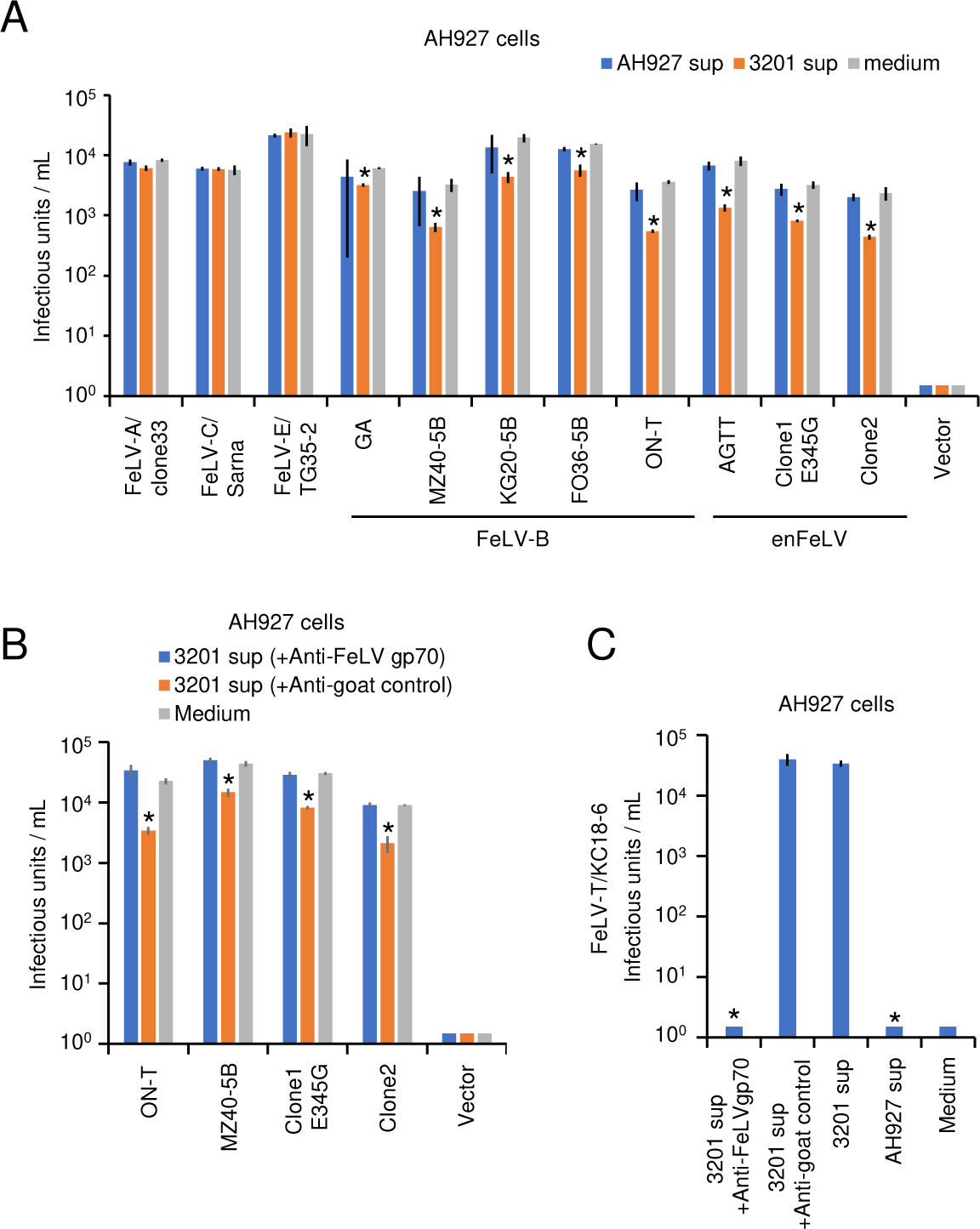
Inhibitory effect of FeLIX from the supernatant of 3201 cells for enFeLV and FeLV-B infection. (A) Inhibition assays of Env-pseudotyped, enFeLV, and FeLV-B viruses were conducted using the culture supernatant from 3201 cells in AH927 cells as target cells. (B) The culture supernatants from the 3201 cells were treated with either goat anti-FeLV gp70 antibody or normal goat serum (control), after which the culture supernatant from which FeLIX was removed and used for the inhibition assays for enFeLV and FeLV-B. (C) The culture supernatant from the 3201 cells and the culture supernatant from which FeLIX was removed were used for the infection assay for FeLV-T. The infectious units (IU) were determined by counting the number of log_10_- galactosidase (LacZ)-positive cells per milliliter (mL) of virus (x-axis). Virus infection titers with standard deviations represent the means of three independent experiments. Medium represents the negative control. Comparisons were performed using Student’s t- test (\**p* < 0.01).

Next, the supernatant of 3201 cells was subjected to FeLIX depletion using an anti-FeLV gp70 antibody (gp70; National Cancer Institute [NCI], Frederick, MD, USA) to evaluate the specificity of FeLIX. The inhibition of FeLV-B (ON-T and MZ40-5B) and enFeLV (clone1 E345G and clone2) infection disappeared; however, treatment with normal goat antisera did not affect the inhibitory effect of the supernatant from 3201 cells (Figure 5B). Furthermore, FeLV-T infection was monitored to assess whether this antibody depleted FeLIX in the supernatant of the 3201 cells. First, the supernatants from 3201 cells supported FeLV-T infection, whereas the supernatants from AH927 cells, which did not express FeLIX, did not support FeLV-T infection. FeLV-T infection was lost in the FeLIX depletion experiment using an anti-FeLV gp70 antibody (Figure 5C).

### Thermal sensitivity of FeLIX

The inhibitory effect of FeLIX following heat treatment was examined; culture supernatants from 3201 cells and HEK293T cells transfected with FeLIX were treated at 56 °C for 30 min. Results showed that the inhibitory effect of FeLIX was completely abolished at 56 °C for 30 min. Furthermore, the effect of heat treatment of FeLIX on FeLV-T infection as a co-factor was examined. The supernatant of 3201 cells was serially diluted and treated at 56 °C for 30 min to examine FeLV-T infection. Results showed that when FeLIX was present in excess, FeLV-T infection was observed even after heat treatment. However, diluted FeLIX abolished the FeLV-T infection after heat treatment (Figure S10). These results suggest that FeLIX is heat-sensitive, and the antiviral activity and co-factor of FeLIX were abolished at 56 °C for 30 min.

### Inhibitory effect of FeLIX for non-feline mammalian retroviruses

Non-feline retroviruses, GaLV, Koala retrovirus-A (KoRV-A), HPG, and 4070A amphotropic murine leukemia virus (4070A MuLV) have been reported as entry receptors (42-45), and their hosts are monkeys, koala, and bats (*Pteropus alecto*), respectively (44-46). We first assessed whether these viruses could infect feline cells. GaLV, KoRV, HPG, and 4070A MuLV cells were susceptible to AH927 and CRFK cells with high viral titers (Figure 6A). Furthermore, GaLV and HPG can infect both MDTF- fePit1 and MDTF-fePit2, whereas KoRV can only infect MDTF-fePit1. Since 4070A MuLV could infect MDTF cells (Figure 6B), the virus receptor was determined in Chinese hamster ovary (CHO) cells, which are not permissive to 4070A MuLV infection. We found that 4070A MuLV could infect CHO cells expressing fePit2 (CHO-fePit2), but not CHO cells expressing fePit1 (CHO-fePit1; Figure S11A). These results suggest that these non-feline retroviruses have the potential to cause infection via the feline phosphate transporter. GaLV and HPG use both fePit1 and fePit2, KoRV uses fePit1, and 4070A uses fePit2 as their receptors.

**Figure 6.**
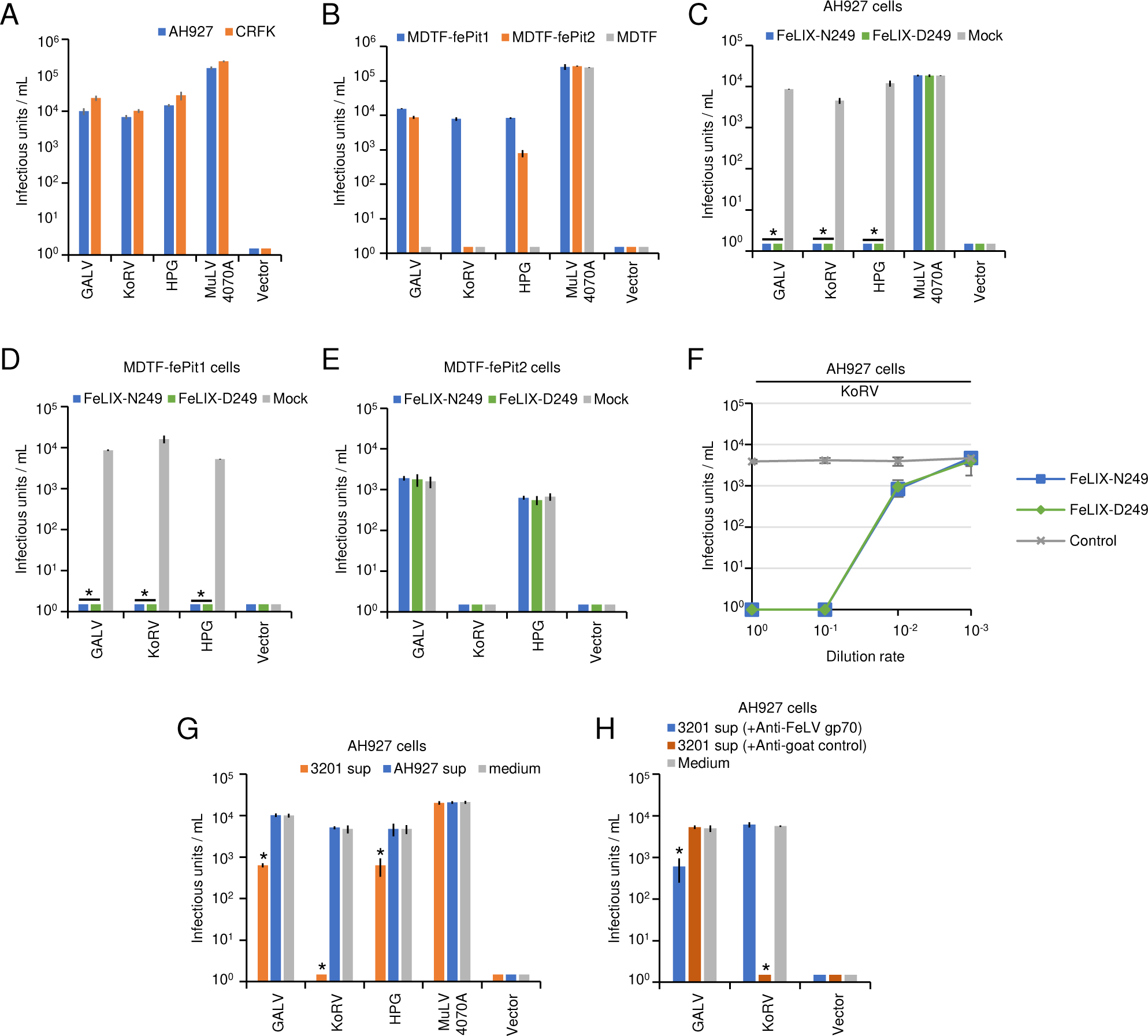
Inhibitory effect of FeLIX for non-feline mammalian retrovirus infection. (A) Infection of non-feline mammalian retroviruses (GaLV, KoRV, HPG, and MuLV 4070A) for feline, AH927, and CRFK cells for receptor usage in (B) MDTF/fePit1,MDTF/fePit2, and MDTF/empty vector. Supernatants from HEK293T cells transfected with expression vectors encoding FeLIX-N249 and FeLIX-D249 were used for inhibition assays against non-feline mammalian retroviruses (GaLV, KoRV, HPG, and MuLV 4070A) in (C) AH927 cells, (D) MDTF-fePit1 cells, and (E) MDTF-fePit2 cells. (F) Dose-dependent inhibition of FeLIX for KoRV Env-pseudotyped viral infection in AH927 cells. (G) Inhibition assays of Env-pseudotyped virus, KoRV, GaLV, HPG, and 4070A ampho-MuLV were conducted using the culture supernatant from 3201 cells in AH927 cells. (H) The culture supernatants from the 3201 cells were treated with either goat anti-FeLV gp70 antibody or normal goat serum (control). Subsequently, the culture supernatant from which FeLIX was removed was used for the inhibition assays for KoRV, GaLV, and HPG infection. The infectious units (IU) were determined by counting the number of log_10_-galactosidase (LacZ)-positive cells per milliliter (mL) of the virus. The viral titers are illustrated as the log number of infectious units (IU) per milliliter (mL). Virus infection titers with standard deviations represent the means of three independent experiments. Mock represents the negative control. Comparisons were performed using Student’s t-test (\**p* < 0.01).

Next, we tested the inhibitory effects of FeLIX-N249, FeLIX-D249, and Trunc-C4 on non-feline retroviruses KoRV-A, GaLV, HPG, and 4070A MuLV. The results showed that they exhibited marked inhibitory effects against viral infections in AH927 and CRFK cells. However, 4070A MuLV was not inhibited by FeLIX-N249, FeLIX-D249, or Trunc-C4 in the feline cells (Figures 6C, S11B, and S12). Further experiments showed that infection with these viruses was inhibited by FeLIX in MDTF-fePit1 cells, but not in MDTF-fePit2 cells (Figure 6D and 6E). A dose-dependent inhibitory effect of FeLIX on KoRV infection was observed (Figure 6F and S11C). The supernatant of 3201 cells significantly (*p <* 0.01) inhibited GaLV, KoRV, and HPG infection in AH927 cells. In particular, the inhibition of KoRV infection by the supernatant of 3201 cells was effective (Figure 6G). The inhibitory effect of the supernatant from 3201 cells on GaLV, KoRV, and HPG infection disappeared after the depletion of FeLIX using the antibody, whereas treatment with normal goat antisera did not affect the inhibitory effect of the supernatant of 3201 cells (Figure 6H). Overall, these results indicated that FeLIX could also confer resistance against non-feline retroviral infections.

## DISCUSSION

Transmission of FeLV-B among domestic cats does not naturally occur, and resistance to FeLV-B infection has been demonstrated in *in vivo* infection models (28). This phenomenon has been suggested because of the presence of enFeLV, and the finding of FeLV-B infection in Florida panthers lacking enFeLV highlights the association between enFeLV and FeLV-B (36). Besides, the presence of ERV is known to provide resistance to exogenous retroviral infection (5-9). However, the molecular mechanisms restricting FeLV-B infection in domestic cats have long remained unknown (37). While FeLIX has been identified as a co-factor for FeLV-T infection, it is unlikely that FeLIX emerged evolutionarily to facilitate viral infection because of its detrimental function in domestic cats. We attempted to address these questions that have been elusive to date. Based on previous findings on identifying antiviral effects by defective Env proteins (5-7), we have proposed plausible explanations for these unresolved issues. In particular, this study highlights that the presence of defective Env is beneficial to the host and that Env, which is scattered throughout the mammalian genome, functions as an antiviral molecule.. Notably, the presence of antiviral mechanisms also highlights the possibility of cross species utilization.

In this study, we determined that FeLIX is a restrictive factor for the feline retroviruses FeLV-B and enFeLV, as well as for the mammalian retroviruses KoRV, GaLV, and HPG. The antiviral machinery is driven by truncated Env proteins encoded by enFeLVs secreted from feline cells. We determined that the antiviral effect of FeLIX is dependent on fePit1. These results implied that receptor interference was the mechanism underlying FeLIX activity. Therefore, FeLIX may function as a barrier to intraspecies and interspecies transmission of retroviruses. These results suggested that truncated Env may constitute an antiviral system in cats. Although soluble Env exhibits antiviral activity, it retains the ability to promote viral infection by FeLV-T, which is detrimental to the host. Antiviral activity may have emerged in an evolutionary arms race, but the ability to promote viral infection may have been created by the emergence of new exogenous retroviruses. Our findings provide insights into the coevolutionary history of retroviruses and their hosts. In particular, we focused on these retroviruses, which infect in a Pit-dependent manner and spread infections.

FeLIX and Trunc-C4 are truncated Env proteins that have an SP and an N-terminal region of SU, which is a putative RBD, but lack the C-terminal region of SU and TM due to a premature stop codon. The amino acid sizes of FeLIX and Trunc-C4 are 273 and 369, respectively. We demonstrated that both molecules exhibit antiviral activity; however, their antiviral activity differed as FeLIX was more effective in inhibiting infection. This highlights the importance of factors other than the structural similarity of the truncated Env. Moreover, FeLIX with a single amino acid substitution between FeLIX-N249 (MGPNP) and FeLIX-D249 (MGPDP) within the FeLV-A neutralization epitope “MGPNL” showed comparable antiviral activities in both AH927 and CRFK feline cells. However, when evaluated in human cells (HEK293T), different activities were observed during enFeLV infection. Amino acid mutations at the neutralization epitope of FeLV-A are present in both FeLIX-N249 and FeLIX-D249, and two enFeLV Env sequences harbored mutations in this epitope. This amino acid mutation may appear to specifically preserve the antiviral effect of FeLIX by neutralizing antibodies. In addition, it has been suggested that viruses with the MGPNL epitope have an *in vivo* growth advantage relative to FeLV-A (47). Thus, this amino acid change may have emerged as an adaptive response during molecular coevolution. The truncated Env protein lacking TM is not tied to the cell membrane and is thus efficiently released from the cells (48). Thus, the RBD plays a crucial role in facilitating the initial stage of the interaction between the viral envelope and its receptors, enabling viral entry (41, 48, 49). Earlier reports have shown that the expression of *env* genes in ERVs can prevent viral infection through receptor interference mechanisms (e.g., Fv-4, Rmcf, Rmcf2, Refrex-1, and Suppressyn) (6-9). Refrex-1 has a structure similar to FeLIX, as it contains the RBD and is primarily secreted in a soluble form, which can offer protection to cells that express both proteins as well as to cells that do not express them. Furthermore, the investigation of enFeLV-derived FeLIX and its evolution revealed that it uses Pit1 as the entry receptor, clearly demonstrating the mechanism underlying the antiviral effect of FeLIX. These findings clearly suggest that truncated Env exerts antiviral effects through competitive receptor binding.

Previous studies reported that the culture supernatant of FeLV-negative, feline 3201 cells contained FeLIX as a co-factor for FeLV-T infection (16). Contrary to its role in supporting viral infection, the present study showed that FeLIX in the culture supernatant of 3201 cells inhibited retroviral infection (enFeLV, FeLV-B, KoRV, GaLV, and HPG). There have been limited reports regarding the effect of FeLIX on FeLV-B infection; however, most of them used FeLIX to investigate the cofactors of FeLV-T infection (16, 33). This is a crucial difference from previous reports. A previous study reported FeLIX in cat serum as a co-factor of FeLV-T infection, whereas FeLV-B only served as a control for infection (33) and lacked a specific means for evaluating the inhibition assay, and a separate study suggested that FeLV-B RBD could inhibit FeLV-B infection (41). Moreover, in our study, feline cells were used for the assay, and the protocol involved incubating the truncated Env proteins prior to retrovirus infection. Additionally, two functions of FeLIX were found to be inactivated by treatment at 56 °C for 30 min under conditions that inactivate complement (Figure S10). Heat-inactivation is required to inactivate the complement before conducting experiments (50). Therefore, the antiviral activity of the serum could not be evaluated. Notably, although small amounts of FeLIX facilitate viral infections, the amount of FeLIX required to block viral infections is 50 times higher (Figure S11). The biochemical activity required for viral infection differed from that involved in inhibiting viral infection. The antiviral effect of the 3201 cell supernatant was not effective against extremely high-titer viruses (Figure S9). Our study also revealed that the antiviral effect varied among FeLV-B strains, with the FeLV-B/GA strain being the least susceptible to the supernatant of 3201 cells. These factors may be reasons why the antiviral effect of FeLIX has not been addressed.

This study revealed that enFeLV uses only fePit1, instead of fePit2, as an entry receptor by analyzing the amino acid mutation of enFeLV Env and receptor usage. We found that FeLV-B typically uses fePit1 as a receptor; however, some FeLV-B strains can use both fePit1 and fePit2. Our findings reveal that, unlike observations from previous studies (17), FeLV-B/GA primarily infects fePit1 but not Pit1 and Pit2. Furthermore, a mutant amino acid change at position 73 (arginine) was responsible for enhanced binding to the Pit2 receptor, consistent with the results of a previous study (17). In addition, the amino acid of lysine at position 66 can affect receptor usage through FeLV-B expansion to Pit2. Gene sequences showing lysine and arginine at positions 66 and 73, respectively, in FeLV-B Env were not found in any enFeLV clones or in the database, suggesting that these amino acid sequences are not derived from the enFeLV *env* gene sequence but may be acquired by genetic mutation. These findings suggest that shifting receptors may have occurred to escape host selection pressure, such as the presence of FeLIX, enFeLV, or the host immunity response. Similarly, FeLV-B showing Env structure diversity may have occurred through host selection pressure, as FeLV-B is a recombinant virus that consists of FeLV-A and an enFeLV *env* gene (27). This seems to be consistent with other studies that revealed a recombinant virus between FeLV-A and the ERV-DC *env* gene in FeLV- D (51); it also occurred on RD-114 by recombination of ERV-DC and a BaEV-related *env* gene (52-55) in response to the selection pressure of the host defense. Selection pressure could trigger the generation of viral variants that avoid these barriers; however, in a positive sense, ERV provides a naturally essential antiviral defense and immunity to the host (5-9). As a result, we believe that an ERV, such as the truncated Env, plays an essential evolutionary role against retroviral infections. We speculate that enFeLV-truncated Env plays a key role in the selective recombination of FeLV-A and enFeLV to generate FeLV-B in domestic cats. The presence of FeLIX in cats could affect the frequency of FeLV-B transmission and the severity of the disease caused by FeLV-B. In particular, it can affect the risk of transmission from domestic cats to other animals. This is consistent with previous studies suggesting that the noncoding RNA of enFeLV-LTRs also tends to control FeLV replication via an RNA interference mechanism (38). Additionally, another study suggested that FeLV-B Env confers strong resistance against feline SERINC5 that depends on its GlycoGag contributing to resistance (56). These results explain a possible antiviral synergistic phenomenon in cats regarding the role of ERV in protecting against the FeLV infection.

Wildlife retroviral models include KoRV, GaLV, and HPG. KoRV is found in koalas and causes serious illnesses. KoRV appears to endogenize and may contain env-ERV in its genome to protect against viral infections (57). In Australia and Asia, HPG was first isolated from the bat genome (*Pteropus alecto*) (45). *Pteropus Alecto* has a large flying range, which increases the possibility of viruses spreading throughout the area and may lead to disease outbreaks in various species. KoRV, GaLV, and HPG have been reported to use Pit1 as receptors (42, 43, 45). Our findings demonstrate that FeLIX inhibited related-Pit1 tropism in retroviruses such as KoRV, GaLV, and HPG. This implies that in the limitation function of the same virus entry mechanism, truncated Env can potentially act in natural host immunity, not only to protect intraspecies but also to protect interspecies against current viral infections. However, new retroviral variants may emerge due to mutations or recombinants that escape host selection pressure. Here, we emphasize that the truncated Env potentially blocks retroviral infection in a wide range of species by targeting a shared viral entrance receptor.

The prevention of viral infection by soluble Env proteins may lead to the consideration of next-generation therapeutic strategies for infectious diseases, including severe acute respiratory syndrome coronavirus 2 (SARS-CoV-2) infections since soluble angiotensin-converting enzyme 2 (ACE2) decoys are highly effective in blocking infection of all variants of SARS-CoV-2 (58, 59).

Finally, although the antiviral effects of FeLIX and defective Env protein can be demonstrated *in vitro*, the exact roles of these soluble proteins *in vivo* remain unclear, which is a limitation of this study.

In conclusion, our findings provide evidence of an antiviral agent against FeLV-B infection in domestic cats that can be extended to non-feline mammalian retroviruses. Furthermore, it conveys a fundamental understanding that soluble Env proteins restrict retroviruses from diverse host species through binding interactions with a common entryway receptor and may play an essential role in protecting against retroviral infection, implying that their functions help control retroviral spread for host immunity and antiviral defense. This study portrays an evolutionary scenario of host-pathogen interactions and may have significant implications for developing treatments for infectious diseases, including vaccine design.

## MATERIALS AND METHODS

### Samples

Domestic cats (7) and European wildcats (60) samples were described previously. Briefly, a specific-pathogen-free (SPF) cat, a 2-month-old female, was obtained from the Nippon Institute for Biological Science, was euthanized, and an autopsy was performed. The study on European wildcats in Navarra, Spain, was conducted with the support of local authorities. Samples were collected from carcasses between 2000 and 2007 and stored frozen at -18 °C. Tissue samples were collected and sent to the Department of Animal Pathology, Faculty of Veterinary Medicine, University of Zaragoza for necropsy. Animal tissues were stored at -80 °C until DNA or RNA was extracted for further investigation.

### Cell lines

The cells were cultured in Dulbecco’s modified Eagle’s medium (DMEM, FUJIFILM Wako Pure Chemical Corporation, Osaka, Japan) supplemented with 10% fetal calf serum (FCS) and 1X penicillin-streptomycin. Cells were incubated in a CO_2_ incubator at 37 °C. In this study, we used the following cell lines: HEK293T (human embryonic kidney transformed with SV40 large T antigen) (61), MDTF (*Mus dunni* fibroblast tail) (62), CHO (63) AH927 (feline fibroblast cells) (64), CRFK (Crandell-Rees feline kidney) (65), 3201 (feline lymphoma cells) (66), GPLac (an *env*-negative packaging cell line containing murine leukemia virus (MuLV) *gag-pol* gene and beta-galactosidase (LacZ)- coding pMXs retroviral vector) (51), and 293Lac cells containing a LacZ-coding pMXs retroviral vector (67). MDTF cells expressing feline phosphate transporter protein 1 (MDTF-fePit1), MDTF cells expressing feline phosphate transporter protein 2 (MDTF-fePit2), CHO cells expressing feline phosphate transporter protein 1 (CHO-fePit1), and CHO cells expressing feline phosphate transporter protein 2 (CHO-fePit2) were established and cultured in the same medium under the aforementioned conditions.

### Establishment of cell lines expressing feline Pit1 and Pit2

Feline Pit1 and Pit2 plasmids were described in a previous report (17). PLAT-A (amphotropic MuLV) packaging cells were transfected with expression vectors (pMSCVneo-fePit1, pMSCVneo-fePit2, or pMSCVneo empty vector) using the TransIT®-293 reagent (Takara, Kusatsu, Japan). Two days later, the supernatants were collected, filtered through a 0.22-µm filter, and then used to infect MDTF cells in the presence of polybrene (10 g/mL). PLAT-GP (an *env-*negative) packaging cells were co-transfected with MuLV 10A1 *env* gene-expression vector, along with the pMSCVneo-fePit1, pMSCVneo-fePit2, or pMSCVneo empty vector, using TransIT®-293 reagent (Takara). Two days later, the supernatants were collected, filtered through a 0.22-µm filter, and used to infect CHO cells in the presence of polybrene (10 g/mL). The cells were cultured in a medium containing 600 μg/mL neomycin (G418) for 2 weeks. These cells were termed MDTF-fePit1, MDTF-fePit2, MDTF-empty, CHO-fePit1, CHO-fePit2, and CHO-empty for further analysis.

### PCR

KOD-ONE Blue (Toyobo, Japan), KOD FX Neo, KOD plus Neo (Toyobo, Osaka, Japan), and GoTaq (Promega, Madison, WI, USA) polymerases were used according to the manufacturer’s instructions.

### PCR product cloning

PCR products were cloned directly into the pCR4Blunt-TOPO (Invitrogen, Carlsbad, CA, USA) or pUC118 vectors (Mighty Cloning Kit; Takara, Kusatsu, Japan). The intact *env* gene was inserted into the pFUΔss expression vector (51). Sequencing was performed using a Contracted Service (Fasmac Corporation, Atsugi, Japan).

### Cloning of enFeLV proviruses and construction of Env expression vector

The construction of a genomic library using cat DNA has been reported previously (51). Briefly, splenic DNA from a single cat (ON-T) suffering from lymphoma due to FeLV infection was digested with EcoRI and ligated to Lambda DASH II/EcoRI vectors (Agilent Technologies, Santa Clara, CA, USA). DNA libraries were screened using a DIG-labeled enFeLV LTR probe generated by PCR with primers Fe-36S and Fe-60R in feline normal chromosomal DNA. The PCR DIG Probe Synthesis kit for probe synthesis and the CSPD luminescent detection kit for visualization of the bound probe were used according to the manufacturer’s instructions (Roche Molecular Biochemicals, Mannheim, Germany). The flanking genomic sequences of the enFeLV proviruses were determined from positive phages, and full-length enFeLV proviruses were amplified from normal cat DNA or ON-T chromosomal DNA using specific primers (Table S1). The amplicons were cloned into pCR4 blunt-TOPO (Invitrogen, Carlsbad, CA, USA) and sequenced (Fasmac Corporation, Atsugi, Japan).

Env expression vectors of enFeLV-clone1, clone2, and clone3 were constructed through PCR amplified using primes pair Fe-560S and Fe-550R based on the enFeLV proviruses in TOPO vector and were cloned into pFUΔss between the BamHI and EcoRI restriction sites (Table S1). enFeLV-AGTT derived from enFeLV-clone3 was constructed by site-directed mutagenesis using the primer pair Fe-661S and Fe-642R. Env expression vectors were constructed using KOD-ONE Blue (Toyobo, Osaka, Japan) following the manufacturer’s protocol. The resulting Env expression plasmids were confirmed through sequencing (Fasmac Corporation, Atsugi, Japan).

### Construction of truncated Env expression vectors

The expression vector pFUΔss was used to construct the expression plasmids. The genes were PCR-amplified from each plasmid using specific primers with enzyme sites, and the PCR products were digested with each restriction enzyme and cloned into the pFUΔss expression plasmid. The Env expression plasmid for pFUΔss FeLIX-D249 was constructed via site-directed mutagenesis using the primer pair SDM-FeLIX1 and SDM- FeLIX2 based on the pFUΔss FeLIX-D249 expression plasmid (7). enFeLV-clone4 (referred to as Trunc-C4) was constructed with primer pairs enFeLVc4-F1 and Fe-686R and FeLV-B/GA RBD (FeB-RBD) with RBD-F1 and Fe-716R and then cloned into the pFUΔss between the *BamH*I and *EcoR*I restriction sites (Table S1). The resulting mutants and Env expression plasmids were confirmed using sequencing (Fasmac Corporation, Atsugi, Japan).

### Construction of chimeric replication-competent virus

Replication-competent FeLV-A/61E carrying the enFeLV-AGTT *env* gene were constructed by replacing the *env* gene. Briefly, the *env* gene of enFeLV-AGTT was PCR-amplified using primers Fe-721S and Fe-748R. The amplicon and FeLV-A/61E provirus digested with SnaB1 and Sph1 (Takara, Kusatsu, Japan) were ligated using In-Fusion HD Cloning Kits according to the manufacturer’s protocol (Takara). The resulting chimeras, termed 61E/AGTT, were confirmed using sequencing (Fasmac Corporation, Atsugi, Japan). 293Lac cells were transfected with 61E/AGTT using the TransIT®-293 reagent. The culture supernatant was confirmed to infect AH927 and CRFK cells at high titers using LacZ staining. To prepare replication-competent viruses, 293Lac cells transfected with FeLV-A/61E, FeLV-B/Gardner–Arnstein (GA), or 61E/AGTT were cultured, and the supernatants were used as the virus stock. The supernatant from HEK293T cells persistently infected with FeLV-B/ON-T (51) was used to infect AH927 cells carrying LacZ (AH927Lac cells). The cell supernatant was used as viral stock. Notably, the supernatant of AH927Lac cells infected with FeLV-B ON-T cells contained FeLV-B and replication-defective FeLV-D, with a low viral titer.

### Construction of Env mutants

Env mutants from enFeLV-clone1 were compared to enFeLV-clone2, enFeLV-clone3, and enFeLV-AGTT to identify amino acid candidates for Env mutants. Mutants derived from enFeLV-clone 1 were constructed by site-directed mutagenesis using the plasmids enFeLV-clone1, E345G, and N394K. The primer pairs used for site-directed mutagenesis were enFeLV-clone1 E345G (Fe-577S and Fe-569R) and N394K (Fe-578S and Fe- 570R). Env mutants derived from FeLV-B/GA, FeLV-B/FO36-5B, and FeLV-B/ON-T were constructed via site-directed mutagenesis using the following plasmids (primer pairs): FeLV-B/GA Q73R (Fe-761S and Fe-788R), FeLV-B/FO36-5B K66D (Fe-782S and Fe-807R), and FeLV-B/ON-T R73Q (Fe-792S and Fe-812R). The mutants were constructed using KOD-ONE Blue (Toyobo, Osaka, Japan) following the manufacturer’s protocol. The resulting mutants were confirmed by sequencing.

### Env-pseudotyped virus preparation

The preparation of Env-pseudotyped viruses carrying LacZ as a marker has been described previously (51). Briefly, GPLac cells were seeded at a concentration of approximately 1 × 10^6^ cells in a six-well plate 1 day prior to transfection. The Env expression plasmids used for viral preparation were as follows: pFUΔss FeLV-B/GA (FeLV-B Gardner–Arnstein Env), pFUΔss FeLV-B/MZ40-5B Env, FeLV-B/KG20-5B Env, FeLV-B/FO36-5B Env (26), FeLV-B/ON-T Env, FeLV-B mutants (FeLV-B/Q73R Env, FeLV-B/ FO36-5B K66D Env, FeLV-B/ON-T R73Q Env, FeLV-A/clone33 Env (68), pFUΔss FeLV-C/Sarma (69), pFUΔss FeLV-E/TG35-2 (70), pFUΔss FeLV-T/KC18-6 (26), enFeLV-clone1 E345G Env, enFeLV-clone2 Env, enFeLV-clone3 Env, enFeLV-AGTT, GaLV Env, KoRV Env (opt-KoRV-A2), and HPG Env (opt-HPG). The mammalian retrovirus *env* genes (opt-KoRV-A2 env and opt-HPG env) were synthesized (Eurofins Genomics, Tokyo, Japan), as shown in Figure S11. The KoRV Env and HPG Env genes were myc-tagged at the C-terminus and inserted into the pFUΔss vector. MuLV 10A1 and MuLV 4070A were purchased commercially (Novus Biologicals). GaLV Env was PCR-amplified from PG13 cells (71) and cloned to the pFUΔss vector. The culture supernatants were collected 48 h after transfection (TransIT-293 transfection reagent), filtered through a 0.22-µm filter or centrifuged at 15,400 × *g* for 2 min at 4 °C, and stored at -80 °C as virus stock for further experiments.

### Infection assay

Target cells (approximately 3 × 10^5^ cells/well) were seeded in 24-well plates 1 day prior to infection. Target cells were infected with each 250 µL of Env-pseudotyped virus in the presence of 10 µg/mL of Polybrene (Nacalai Tesque, Kyoto, Japan) for 2 h. After the addition of fresh medium, the cells were cultured for 2 days post-infection. After 48 h of incubation, the cell supernatants were removed, and the cells were fixed with 250 μL of 2% glutaraldehyde for 15 min at room temperature (20–25 °C, stained with 5-bromo-4- chloro-3-indolyl-β-D-galactopyranoside (X-Gal), and counted as blue-stained nuclei, as visualized under a microscope. Viral titers are presented as infectious units (IU) per milliliter with standard deviations.

### Viral interference assay

The viral interference assay was performed as described previously (16, 17). AH927 cells infected with the replication-competent viruses, FeLV-B/GA, and FeLV-B/ON-T, were cultured as AH927/FeLV-B/GA and AH927/FeLV-B/ON-T cells for the interference assay. AH927/FeLV-B/ON-T cells contain replication-defective FeLV-D. Target cells were infected with 250 µL of Env-pseudotyped virus for 2 days in the presence of 10 µg/mL of Polybrene (Nacalai Tesque). Two days post-infection, the cells were stained with X-Gal staining solution by counting the number of blue-stained nuclei, and single-cycle infectivity was measured and visualized under a microscope. Viral titers are presented as IU per milliliter with standard deviations.

### Viral inhibition assay in the presence of truncated Env proteins

HEK293T cells were seeded at a concentration of approximately 1 × 10^6^ cells in a six-well plate 1 day prior to transfection with truncated Env expression plasmids. The supernatants of HEK293T cells transfected with pFUΔss FeLIX-N249, pFUΔss FeLIX- D249, pFUΔss FeLV-B/GA RBD (hereafter termed as FeB-RBD), pFUΔss Env clone4 (Trunc-C4), and pFUΔss empty vector (negative control) for approximately 48 h were the source of truncated Env proteins. The infection assay was conducted as described previously. In 24-well plates, target cells were treated with 250 µL of cell supernatant (source of truncated Env protein) for 2 h. Subsequently, 250 µL of Env-pseudotyped viruses or replication-competent viruses were inoculated into the target cells with 10 µg/mL polybrene for 2 h, after which 250 µL of fresh medium was added. Two days post-infection, the cells were stained with X-Gal. Single-cycle infectivity was determined by counting blue-stained nuclei under a microscope. IU per milliliter and standard deviations were used to represent the viral titers.

### Viral infection assay in the presence of supernatants of cell cultures

A viral infection assay was performed in the presence of cell supernatants from 3201 cells to investigate the inhibitory effects of cell supernatants on viral infection. The culture supernatant of cells for 2 days at a concentration of approximately 3 × 10^6^ cells was collected and kept at -80 °C after it was filtered through a 0.22-µm filter. The viral interference assay was essentially followed by an infection assay in 24-well plates. First, 250 µL of cell supernatant was incubated for 2 h. Next, 250 µL of Env-pseudotyped viruses were inoculated into AH927 cells in the presence of 10 µg/mL of Polybrene (Nacalai Tesque). The supernatants of 3201 cells were depleted using anti-FeLV gp70 SU (gp70; National Cancer Institute [NCI], Frederick, MD, USA). The supernatant was treated for 4 h at 4 °C with goat anti-FeLV gp70 antibody or normal goat serum as a control (Wako Pure Chemical Industries, Ltd., Osaka, Japan), conjugated with protein G-agarose (Santa Cruz Biotechnology), and centrifuged at 15,400 × g for 2 min at 4 °C.

### Detection of fePit1 and fePit2 expression levels via RT-qPCR

Total RNA was extracted from the tissues of a specific pathogen-free cat (Kyoto-SPF1) described in our previous study (7), as well as from the feline cell lines AH927 (feline embryonic fibroblast cells), CRFK (feline kidney cells), Fet-J (PBMC), MCC (feline large granular lymphoma) (72), 3201 (feline T-cell lymphoma), and MS4 (feline B-cell lymphoma) (73) using an RNAiso Plus kit (Takara), in accordance with the manufacturer’s instructions. Thereafter, cDNA was synthesized using a PrimeScript II first-strand cDNA synthesis kit (Takara) according to the manufacturer’s instructions. Prior to reverse transcription, the RNA samples were treated with recombinant DNase I (TaKaRa). The cDNA was amplified using SYBR Premix Ex Taq II (Tli RNase H Plus; Takara) in a CFX96 Touch real-time PCR detection system (Bio-Rad, Hercules, CA, USA). Feline Pit1 was amplified using the primers Fe-844S and Fe-874R. Feline Pit2 was amplified using the primers Fe-845S, Fe-875R, and the internal control feline peptidyl prolyl isomerase A (PPIA) was amplified using the primers Fe-227S and Fe- 204R. Thermal cycling was performed according to the manufacturer’s instructions.

### Detection of FeLIX expression level via qRT-PCR

Briefly, the same cDNA samples were used for the detection of FeLIX, the cDNA tissues of a specific pathogen-free cat (Kyoto-SPF1), and the feline cell lines AH927, CRFK, Fet-J, MCC, MS4, and 3201. The cDNA was amplified using Premix Ex Taq (Probe qPCR; Takara) in a CFX96 Touch real-time PCR detection system (Bio-Rad, Hercules, CA, USA). FeLIX was amplified using primers FeLIX-F and FeLIX-R and the amplification was detected using the probe FeLIX-P (containing 6-carboxy-fluorescein; FAM), and the internal control feline peptidyl prolyl isomerase A (PPIA) was amplified using primers Fe-227S and Fe-204R via SYBR Premix Ex Taq II (Tli RNaseH Plus; Takara). Thermal cycling was performed according to the manufacturer’s instructions.

### Immunoprecipitation and immunoblotting

Plasmids were introduced into HEK293T or GPLac cells using the TransIT®-293 (Mirus) reagent in six-well plates. Two days after transfection, cell pellets were collected by washing with PBS three times. Cells were resuspended in lysis buffer (20 mM Tris-HCl [pH 7.5], 150 mM NaCl, 10% glycerol, 1% Triton X-100, 2 mM EDTA, 1 mM Na_3_VO_4_, and 1 µg/ml each of aprotinin and leupeptin) to prepare cell lysates, which were then incubated for 20 min on ice. Top-speed centrifugation for 20 min at 4 °C was used to remove the insoluble components, and a protein assay kit (Bio-Rad Laboratories, Carlsbad, CA) was used to calculate the protein concentrations.

Cell supernatants were filtered through a 0.22-µm filter and were incubated overnight at 4 °C with the desired antibody. The purified protein (cell lysate and supernatant) was mixed with Sample Buffer Solution (Nacalai Tesque) and then heated at 95–100 °C for 5 min. SDS-PAGE was conducted using 4–20% gels (Invitrogen, Carlsbad, CA, USA) at 100 V for 2 h and then transferred to nitrocellulose filters for western blotting with the following primary antibodies used in the assays: anti-Myc monoclonal antibody conjugated with horseradish peroxidase (FUJIFILM Wako Pure Chemical Corporation; dilution 1:1000), goat polyclonal anti-FeLV gp70 SU (National Cancer Institute [NCI], Frederick, MD, USA; dilution 1:15000), and mouse anti-FeLV TM protein (p15E; antibody PF6J2A; Custom Monoclonals International, CA, USA; dilution 1: 5000). The secondary antibodies were horseradish peroxidase (HRP)-conjugated anti-mouse IgG (Cell Signaling Technology, Danvers, MA; dilution 1:3000) or HRP-conjugated donkey anti-goat IgG (Santa Cruz Biotechnology; 1:10000). The substrate was LumiGLO® Reagent (20×) and 20X peroxide (Cell Signaling Technology in Danvers, Massachusetts). For imaging, the blots were performed using Amersham ImageQuant 800 (Cytiva, Shinjuku, Japan).

### Phylogenetic and sequence analyses

To identify enFeLV-related sequences in domestic cats (*Felis catus*), the *env* gene sequence of enFeLV-AGTT was first BLASTed against all domestic cat genomes available in NCBI (GCA_018350175.1, GCA_016509815.2, GCA_000181335.6, GCA_000003115.1, and GCA_013340865.1). Next, 10 kbp upstream and downstream genomic regions for enFeLV-like viral *env* genes were downloaded under the condition that only proviruses containing an *env* gene were analyzed. Sequences were compared using SEAVIEW (74), BioEdit (75), and Genetyx software (Genetyx Corporation, Tokyo, Japan). All enFeLV-related viruses are listed in Table S2. A phylogenetic tree was constructed using the sequences listed in Table S2. The MEGA11 software package was used for the phylogenetic analysis (76). Alignment was performed using MUSCLE (77). A phylogenetic tree was constructed using the maximum likelihood (ML) method, with robustness evaluated by bootstrapping (1,000 times). The substitution models were selected based on the lowest BIC score (HKY + G) for the LTR (76, 78).

### Estimation of the integration timings of FeLV truncated Env and full-length Env

The NCBI RefSeq genomic database was scanned for enFeLV-related viral env genes by BLAST-ing. The enFeLV provirus sequences were extracted. The estimated integration timing based on the substitution rate of 5’ LTR and 3’ LTR sequences was calculated (Table S3). The mean divergence rate of noncoding regions in the domestic cat genome (1.2 × 10^-8^ substitutions/site/year) was used because the mutation rate of LTRs in enFeLVs is unknown (79). The 5’ LTR and 3’ LTR sequences of each enFeLV were multiple-aligned, and the genetic distance (D) was calculated using the Kimura two-parameter model (80) in MEGA11 software (76).

### Statistical analysis

Data are represented as the mean with standard deviation (SD) in all bar diagrams. The results of assays were considered statistically significant if *p-*values were < 0.05 by Student’s t-test.

## Acknowledgments

This study was funded by the Japan Society for the Promotion of Science KAKENHI (grant number: 20H03152 and 23H02393) to K.N. The funders had no role in study design, data collection and interpretation, or the decision to submit the work for publication.

We are grateful to Hajime Tsujimoto for providing the FeLV-B/pFGB and FeLV-C/pFSC plasmids, Yoshinao Kubo for MDTF cells, Edward Hoover for FeLV-A/61E, Toshio Kitamura for the GP cells and pMxs retroviral vector, and Julie Overbaugh for the feline Pit1 and Pit2 expression plasmids.

## Author contributions statement

The authors declare no conflicts of interest. The contributions of the authors are described as follows. Conceptualization: K.N., Data Curation: D.P., A.M, K.N., Formal analysis: D.P., A.M., K.N., Funding acquisition: K.N., Investigation: D.P., D.T., M.K., L.A., D.M., A.M., K.N. Methodology: D.P., A.M., K.N., Project administration: K.N., Supervision: T.K., A.M., K.N., Validation: D.P., J.K., A.M., K.N., Writing-original draft: D.P., K.N., Writing-review & draft: D.P., K.N.

## Data availability

All data in this study are included in the main text and figures, supplementary data. The sequences described in this paper have been deposited in the DDBJ/EMBL/GenBank under the accession numbers LC196053–LC196055 and LC198317-LC198319.

